# Dynamics-enhanced Molecular Property Prediction Guided by Deep Learning

**DOI:** 10.1101/2025.11.07.687115

**Authors:** Qiang Liu, Debby D. Wang, Weiqing Guo, Yuting Huang, Xizhao Wang

## Abstract

Molecular property prediction (MPP) is a key challenge in computational biology and drug discovery. Traditional approaches mostly rely on feature representations yielded from static structures of molecules, ignoring their dynamic nature. The scarcity of dynamics data in public databases and the complexity of learning such high-dimensional data made it more difficult for dynamics-involved studies. Accordingly, we built a series of dynamics datasets for MPP tasks by performing comprehensive molecular dynamics (MD) simulations on different molecules. In addition, we proposed a dynamically enhanced molecular representation (DEMR) method with multiple sampling strategies for the dynamics frames. Besides, two deep learning pipelines were employed for mapping DEMR to the molecular properties in various tasks. Our models achieved better performance in different MPP tasks, with a practical guidance in efficient frame selection. This study highlights the significance of integrating MD data into MPP tasks and opens new avenues for structure-based drug design. The generated MD datasets are publicly available in a *Zenodo* repository at https://doi.org/10.5281/zenodo.15788151.

## 1 Introduction

Predicting molecular properties with high accuracy is a fundamental goal in drug discovery, computational chemistry, and chemical safety evaluation [1]. These molecular properties can span a wide spectrum of chemical, biological, and pharmacological characteristics of molecules (Fig. 1(a)). In particular, biological properties such as toxicity, bioactivity and binding affinity play crucial roles in the development of new drugs [2], making the in-silico prediction of these properties an active area of research.

**Fig. 1.**
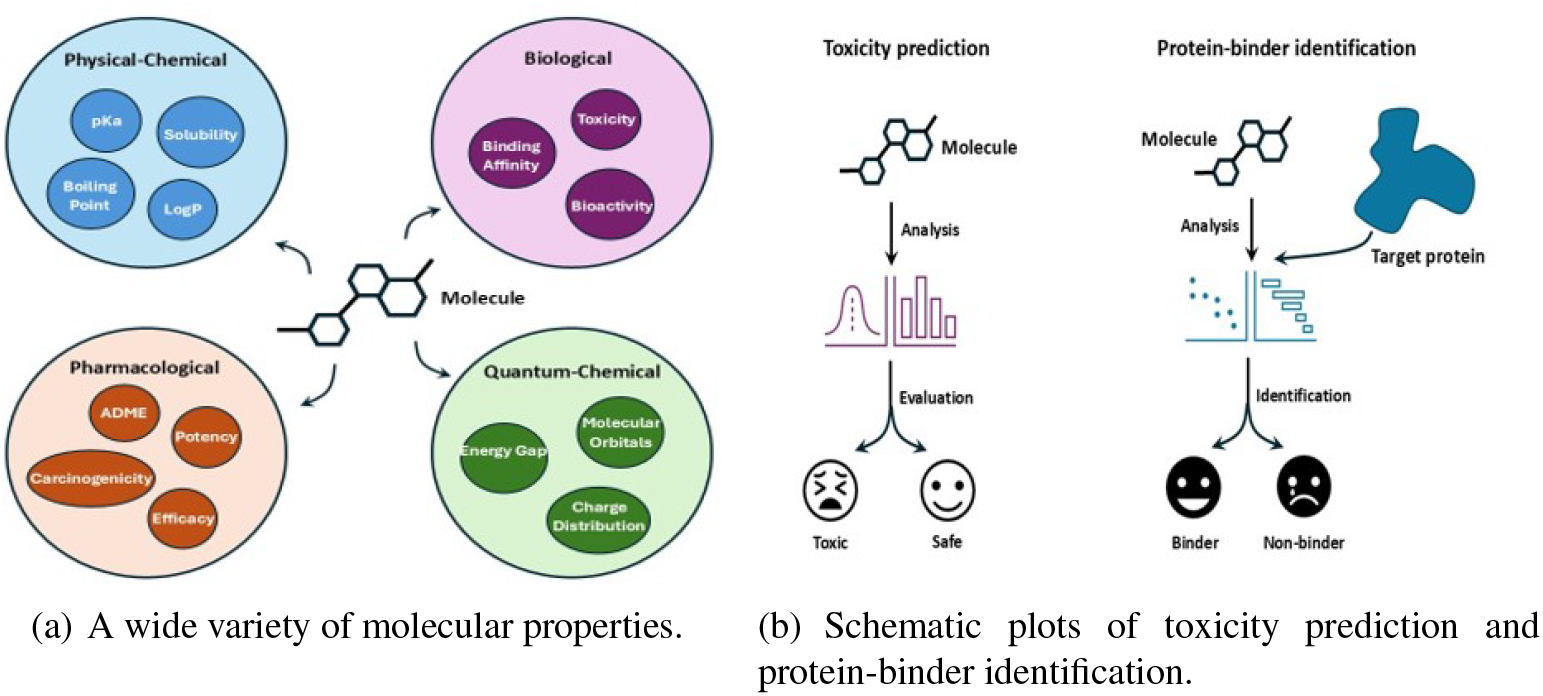
Introduction of molecular property prediction tasks. (a) shows diverse categories of molecular properties spanning chemical, biological, and pharmacological characteristics. (b) shows two representative biological property prediction tasks, including toxicity prediction and protein-binder identification.

*Toxicity prediction* and *protein-binder identification* are two representative tasks in computational drug discovery (Fig. 1(b)). *Toxicity prediction* [3] has the goal of filtering out the compounds with potential safety issues before experimental testing, which helps reduce the failure rate of clinical trials. *Protein-binder identification* [4] aims to recognize small-molecule binders for a target protein, prioritizing them for downstream experimental validation. Aiming at these molecular property prediction (MPP) problems, a variety of computational approaches have been proposed in recent decades, with artificial intelligence (AI) methods emerging as a key focus among them.

During the application of AI techniques, particularly deep learning, to MPP tasks, molecules should be represented as a structured format suitable for computational learning. ***Molecular representation formats***. In earlier studies, a molecule was frequently represented by a series of descriptors, showing its physical and chemical properties (e.g. molecular weight, logP, number of hydrogen-bond donors/acceptors, and polarizability) [5,6,7]. Another popular representation method is molecular fingerprints, such as extended-connectivity fingerprints (ECFPs). These fingerprints often adopt a binary string or a number set to capture the structural patterns of a molecule [8,9,10]. In addition, representing a molecule as a sequence, such as a SMILES or SMARTS string [11,12,13], is another option for subsequent learning. A chemical string encodes molecular structures in a text-based format and is friendly to sequence-oriented learning. In recent years, molecules are often represented as graphs, with atoms treated as nodes and bonds as edges [14,15,16]. This representation method can store the natural structure and topological information of a molecule. ***Learning mechanisms***. Treated as feature vectors, molecular descriptors and fingerprints have been extensively learned by traditional machine learning models (e.g. support vector machines and random forests) [9,17,18] and shallow neural networks (NNs) [19]. For chemical sequence representation, specialized large language models (LLMs) have been developed to learn tokenized sequence features [20,21,22]. A series of graph neural networks (GNNs) have also been devised to handle molecular graphs in MPP tasks [14,15,16,23,24].

Despite the decent predictions, the earlier models mostly treat the input molecules as static. However, molecules are dynamic objects in solution, and their dynamics may drive the bioactivity and functions. Considering molecular dynamics (MD) is therefore an advantage in MPP tasks, but is limited to the lack of dynamics data in public repositories and the difficulty in learning such high-dimensional data. Consequently, this study aims to develop dynamics-enhanced models for MPP, with a focus on the two representative tasks of *toxicity prediction* and *protein-binder identification*. Our main contributions are listed below.

– Comprehensive MD simulations have been conducted on the molecules in several typical MPP sets, facilitating the use of dynamics data by the MPP community.
– The potential of deep learning in handling the high-dimensional dynamics data has been explored, leading to better MPP performance.
– The task-specific sampling frequency for generating MD trajectories has been investigated, in order to further boost the MPP performance.
– The impact of selecting molecular conformations in different RMSD ranges on the MPP tasks has been investigated, which can identify more representative conformations in dynamics-involved prediction works.

## 2 Related Work

In recent decades, AI-based methods have attracted considerable attention in MPP tasks. Early quantitative structureactivity relationship (QSAR) methods relied on manually crafted molecular descriptors/fingerprints (e.g. ECFPs) and were modeled using traditional machine learning algorithms [25]. They remain common baselines for evaluating the performance of MPP models [9,17,18]. With the development of deep learning, a variety of deep neural networks have been used for MPP tasks. Mayr et al. [26] proposed a deep neural network model (*DeepTOX*) for learning molecular fingerprints in MPP tasks, which achieved state-of-the-art performance at the time of its introduction. Convolutional Neural Networks (CNNs) are often used for image processing, but they have recently been introduced to molecular modeling and representation learning. The typical process involves encoding the sequence representation of molecules (e.g. SMILES strings) and applying convolutional operations to the extraction of underlying features [27,28]. In addition, the SMILES string of a molecule can be viewed as a form of chemical language, and many sequence-based models such as the Transformer architecture can be used for learning representations of such chemical languages [29]. With the emergence and rapid development of LLMs, many attempts have been made to use pre-trained LLMs (e.g. *ChemBERTa*) for MPP. LLMs can recognize patterns in molecular data and have shown good predictive performance in a group of benchmark MPP tasks [20,21,22]. Besides sequence representations, graph representations that capture molecular topological information are also popular in MPP tasks.

Molecules can be naturally represented as graph structures, where nodes correspond to atoms and edges represent chemical bonds between them. GNNs can effectively capture the chemical structure and topological connectivity of molecules, demonstrating significant advantages in MPP tasks. Therefore, GNNs are well-suited to learning molecular graph representations [30]. Typical GNN architectures include GCN [31], GIN [32], GAT [33],

GraphSAGE [34], and APPNP [35], and they encode molecular structural features via convolution, message-passing, attention mechanisms, inductive aggregation, and neural propagation, respectively. These models typically learn from 2D topology graphs of molecules, which can be constructed from SMILES using RDKit and other tools [36]. In particular, a series of MPP models have been developed using these GNN architectures, addressing tasks such as *protein-binder identification* [37], *toxicity prediction* [16,38,39], *binding affinity prediction* [14,15] and more [40].

Moreover, to the best of our knowledge, this study is the first to explore MPP tasks in dynamic molecular systems. We further compare our approach with the aforementioned methods to demonstrate the effectiveness and highlight the novelty of our methods.

## 3 Materials and Methods

### 3.1 Generating MD data for MPP tasks

Before constructing dynamics-enhanced models for MPP tasks, we conducted reliable MD simulations on several widely applied benchmark datasets.

– *TOX21* from [MoleculeNet](https://moleculenet.org/datasets-1) are our base data for toxicity prediction. 12 different toxic effects of thousands of molecules were measured by specifically designed assays, constituting *TOX21*. These assays are either related to nuclear receptor signaling pathways (NR type) or cellular <u>s</u>tress <u>r</u>esponse pathways (SR type), demonstrating potential toxic effects of chemical compounds on different biological systems.
– *ADA17, EGFR* and *HIVPR* from [DUD-E](https://dude.docking.org/) are the original data for protein-binder identification. *ADA17* includes a large group of small molecules that are either binders (actives) or non-binders (decoys) to ADAM Metallopeptidase Domain 17 (ADA17). Similarly, *EGFR* and *HIVPR* indicate the molecules for epidermal growth factor receptor (EGFR) and HIV-1 protease (HIVPR), respectively.

The *GROMACS* [41] software package with *CGenFF* (CHARMM General Force Field, details in Appendix A) was primarily used to generate MD data. To ensure reliable MD simulations, we applied several exclusion criteria to the molecules in the aforementioned datasets. The excluded molecules are ➀ those whose initial structures are of poor quality, ➁ those failing to generate MD-required topological files, and ➂ those containing atoms that cannot be recognized by *CGenFF*. Finally, the filtered datasets for each MPP task are presented in Table 1. To verify the diversity of the filtered datasets, we investigated the average pairwise similarity with respect to the positive (toxic) samples, negative (non-toxic) samples, and all samples in each set. As shown in Fig. 2, both toxic and non-toxic molecules exhibit low intra-class similarity, suggesting high structural heterogeneity within each class. The low inter-class similarity between toxic and non-toxic molecules indicates that the structural patterns distinguishing the two classes do not substantially overlap.

**Table 1.**
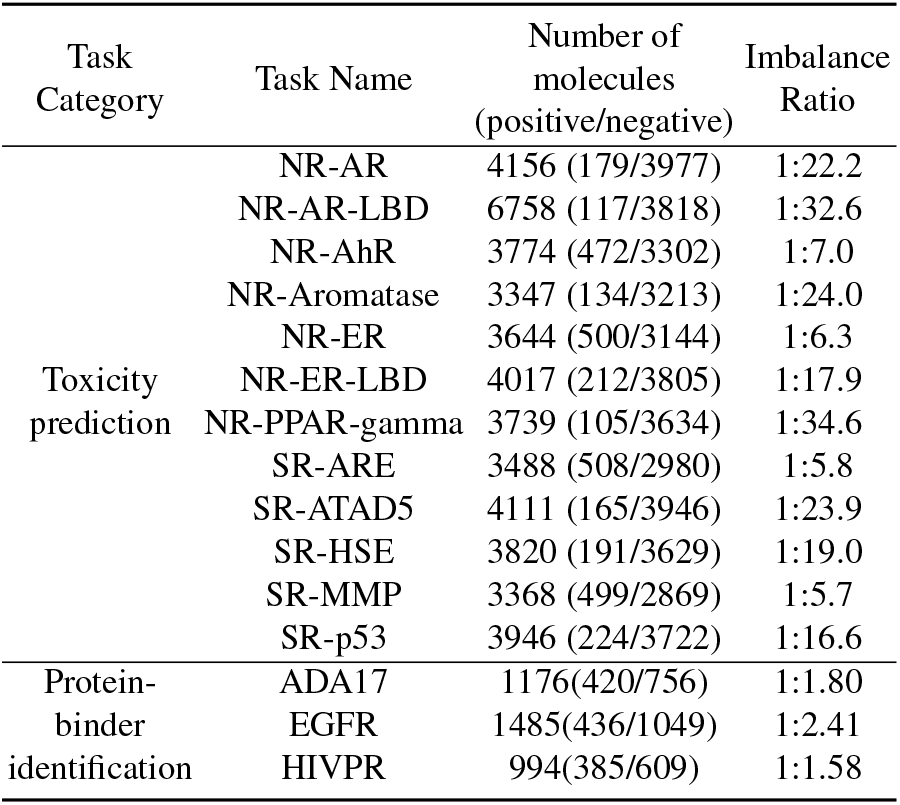
Statistics on Our MD Datasets for MPP tasks.

**Fig. 2.**
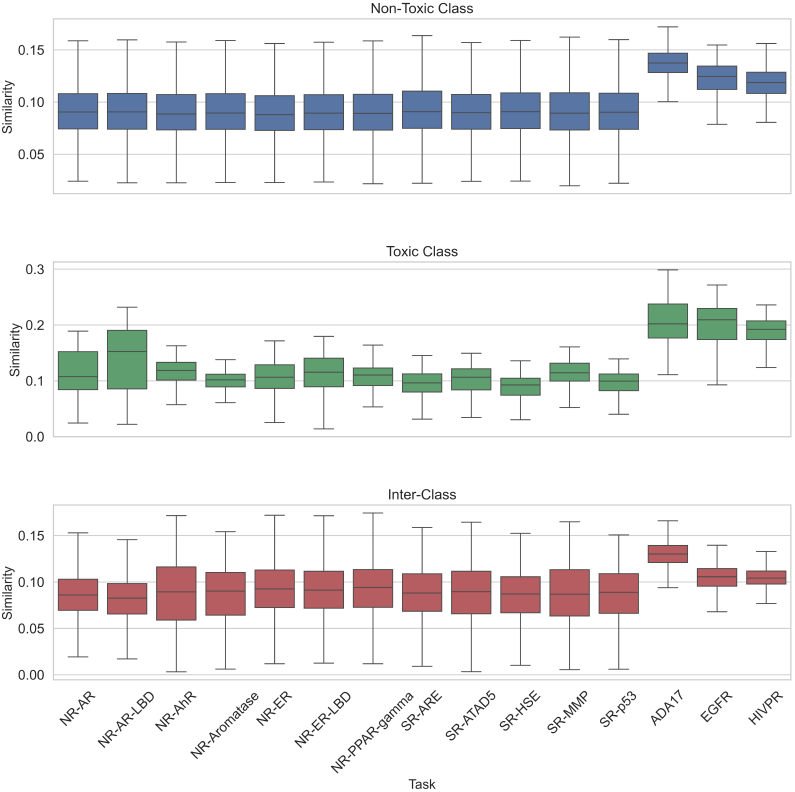
Pairwise similarity for molecules in each filtered dataset.

Based on these datasets, we performed MD simulations on the molecules following the workflow shown in Fig. 3. Implementation details are described below.

1. **Preparation and topology generation**. Generating the 3D structure (in ‘pdb’ or ‘mol2’ format) of a molecule from its SMILES string is the starting point and can be accomplished using the *RDKit* software package. Based on the 3D structure, a molecule can be parameterized using *GROMACS*, and its topological files were generated via the *CGenFF* Web Server.
2. **Setting up the simulation environment**. Since molecules are dynamic in solution, we need to set up a solution environment computationally before the simulations. For each small molecule, a rectangular simulation box was defined, sharing the same center as the molecule and extending at least 1.0*nm* beyond the molecule in all directions. Subsequently, the molecule was computationally solvated, with the *SPC* water model applied. This water environment is the main simulation environment.
3. **Energy Minimization (EM)**. EM is performed to eliminate unfavorable atomic contacts and high-energy conformations. 50,000 minimization steps using the steepest descent algorithm were performed on each system, with the maximum force on the system often reduced to below 1000 kJ/mol/nm. The Particle Mesh Ewald (PME) method was employed to treat long-range electrostatic interactions, with the van der Waals force cutoff distance set to 1.2*nm* and periodic boundary conditions applied in all directions. This procedure ensures a stable initial conformation for the system.
4. **System equilibration**. This process includes both NVT and NPT equilibration phases, which aim to stabilize the system at the target temperature and pressure. *NVT equilibration:* The system temperature was equilibrated to 300*K* using the V-rescale thermostat method for 100*ps* (time step of 0.002*ps* and total 50000 steps) with a leap-frog integrator. Positional restraints were applied to the molecules, and neighbor searching was performed using the Verlet cut-off scheme (1.2*nm* cut-off for non-bonded interactions). *NPT equilibration:* Subsequently, the NPT equilibration for 100*ps* was performed using the Berendsen barostat method to regulate the pressure to 1*bar*, with an isothermal compressibility of 4.5 *×* 10^*−*5^*bar*^*−*1^ and the same positional restraints. The temperature control parameters were kept consistent with those in the NVT phase.
5. **Production MD**. After removing the positional restraints, an all-atom molecular dynamics simulation was performed for 10*ns* using a leap-frog integrator (time step of 2*fs*, totaling 5000000 steps). The system temperature and pressure were maintained. Non-bonded interactions were treated with a cut-off distance of 1.2*nm*. Electrostatic interactions were calculated using the PME method, and van der Waals interactions were corrected with the force-switch algorithm. Periodic boundary conditions were applied in all three spatial dimensions, and dispersion correction was enabled to improve the accuracy of energy and pressure calculations. The MD trajectories were saved every 1*ps* during the simulation for each system.
6. **Quality evaluation**. Due to the use of periodic boundary conditions, molecules may appear ‘broken’ or may ‘jump’ back and forth across the box during the simulation. Therefore, trajectory post-processing was performed to re-center the molecules and restore the integrity of the unit cell. To evaluate the quality of the MD simulations, we examined the temperature (NVT phase), density (NPT phase), total energy (production MD phase), and root-mean-square deviation (RMSD, production MD phase) of each system. The stable trends of these indicators confirm the reliability and convergence of the MD simulation process.

**Fig. 3.**
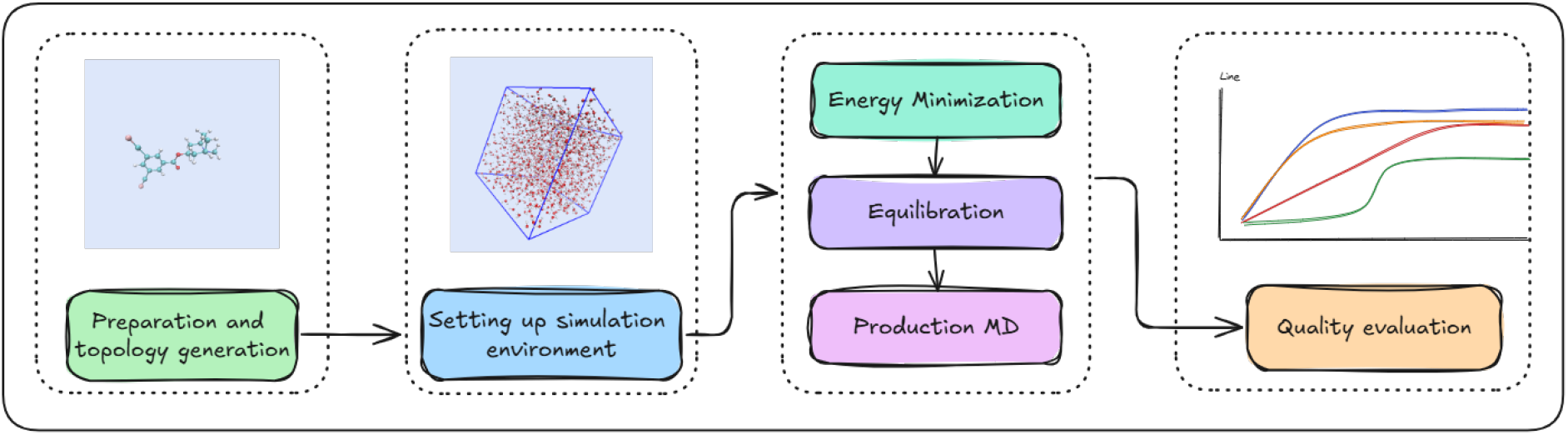
A generally workflow for MD simulations.

### 3.2 Dynamics-enhanced molecular representation modeling

Molecular representations play a crucial role in MPP tasks, and conventional methods often overlook the dynamic nature of molecules. In this study, we introduce a dynamics-enhanced molecular representation (DEMR) method, which describes the structures and dynamics of molecules. Compared with representations relying solely on static molecular structures, DEMR substantially enhances the expressive power and informativeness of molecular features.

Through MD simulations, we collected a 10*ns* dynamics trajectory for each molecule. Such a trajectory contains 10000 frames (structural conformations), and can be denoted as 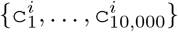 for the *i*-th molecule. A large number of frames were recorded at a relatively short time intervals (1*ps*), because our objective was to examine the impact of the sampling frequency or strategy on the subsequent learning process. Given a molecular trajectory, *representing each frame* 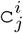 and *sampling the frames further* are the two key points of DEMR.

#### Representing each frame 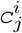

To efficiently capture the structural information of each frame, both the atomic identities and their relative spatial positions are essential. Inspired by Zheng et al. [42], we employed an atomtype-based and distance-guided strategy to represent each molecular frame. Specifically, we defined the following set of atom types.

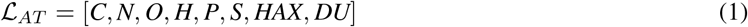

The frame indices *i* and *j* are ignored here for simplicity. *HAX* refers to halogen elements (*F, Cl, Br*, and *I*), and *DU* represents all remaining elements. Further, a list of atom-pair types can be defined as a matrix.

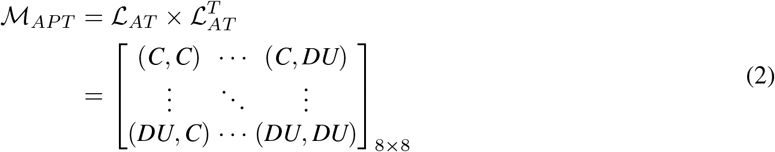

With respect to atomic interactions, we consider the following distance ranges.

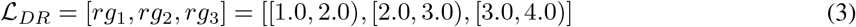

These ranges are defined to encode short-, medium-, and long-range spatial interactions in molecular systems. Specifically, short-range interactions correspond to covalent bonds, medium-range interactions are typically associated with hydrogen bonds and dipole-dipole interactions, and long-range interactions reflect van der Waals forces and hydrophobic contacts [43]. Based on this scheme, a feature representation can be derived for each frame as follows.

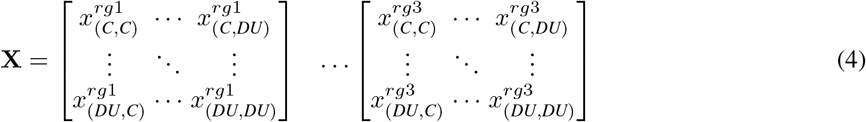

Here 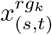 denotes the count of atom pairs of type (*s, t*) in Eq. 2 whose interatomic distance falls within the range *rg*_*k*_. The resulting feature tensor, **X** *∈* ℝ^3*×*8*×*8^, efficiently encodes the key atomic and structural information of the frame.

#### Sampling the frames further

Originally, we collected *J* = 10, 000 frames 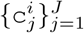 for each molecule, and each frame can be represented as 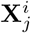 according to Eq. 4. To leverage the dynamic information more efficiently, we construct a **DEMR**^*i*^ tensor as an enriched representation based on different sampling strategies. ➀ Timeinterval sampling. Different intervals along the time axis were used to sample the frames. A series of time intervals *Freq ∈ {*1*ps*, 2*ps*, 5*ps*, 10*ps*, 20*ps*, 50*ps*, 100*ps*, 200*ps*, 500*ps*, 1000*ps}* were adopted in this study, and 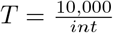 frames were sampled. ➁ RMSD-based sampling. Frames were ranked according to their RMSD values relative to the original conformation and categorized into high-, medium-, and low-level changes (HLC, MLC, and LLC). Focusing on a category (HLC, MLC or LLC), *T* frames were sampled, with *T* selected from *{* 2, 5, 10, 20, 50, 100, 1000*}*. These sampled frames were weighted by their RMSD values during the subsequent learning process. This strategy was designed to evaluate the effect of molecular conformational variation on model performance. ➂ Hybrid-category sampling. To jointly consider all three categories (HLC, MLC, and LLC), 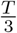 frames were sampled from each category and combined into a set of *T* frames. Learnable weights were then assigned to these frames in the following learning stage to assess the relative importance of different conformational categories for the MPP tasks. Finally, the newly sampled *T* frames yield an enriched representation of the *i*-th molecule as follows:

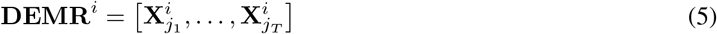

This tensor captures both the static structural features and the dynamic conformational changes of the molecule, providing richer information for developing learning models in MPP tasks.

### 3.3 Dynamics-involved learning architecture design

To effectively learn the **DEMR** tensors, we design several hybrid neural network architectures, as illustrated in Fig. 4.

**Fig. 4.**
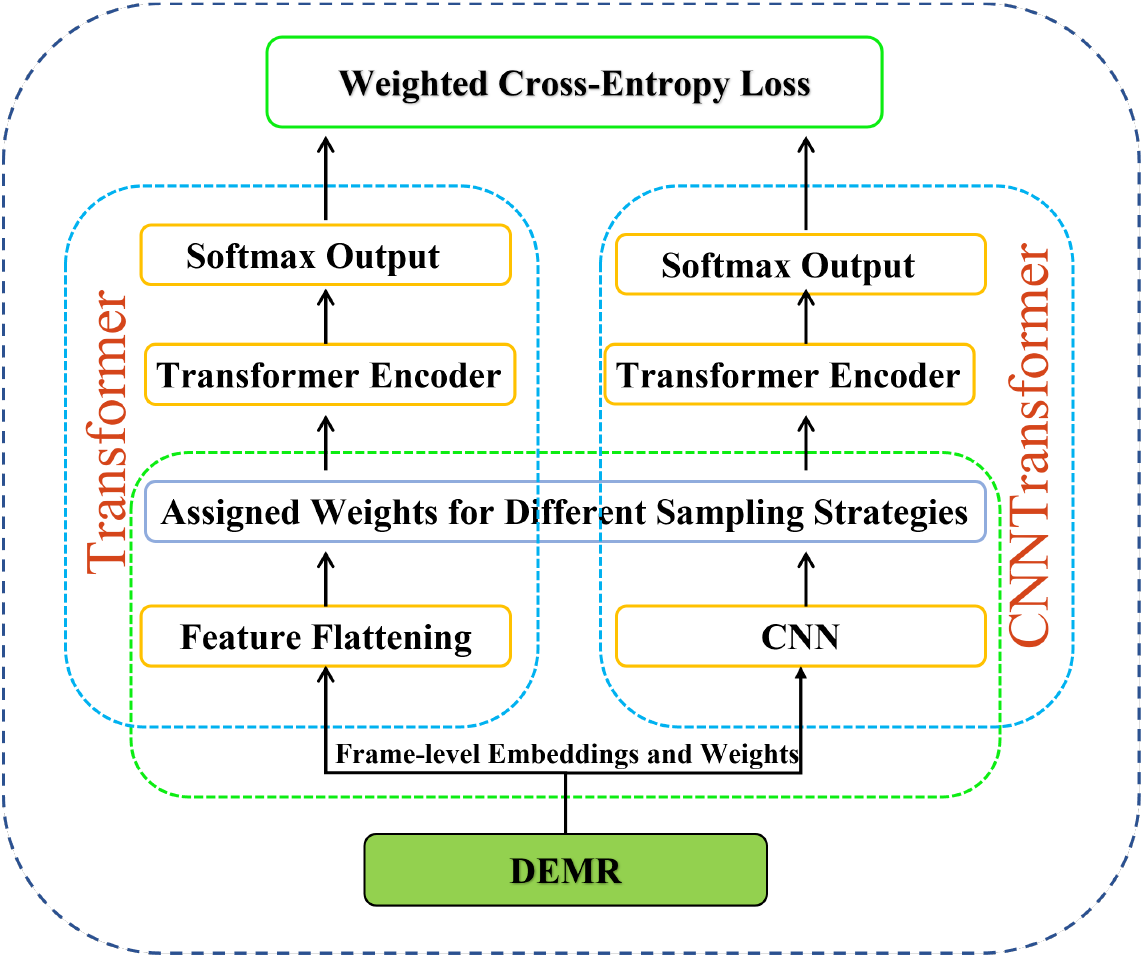
The network architecture for dynamics-enhanced MPP tasks.

#### Frame-level embeddings and weights

The input tensor **DEMR** *∈* ℝ^*T ×*3*×*8*×*8^ consists of *T* structural frames with each frame encoding the co-occurrence of atom type pairs across three distance intervals. The sample index *i* is omitted now for simplicity. For each frame, we map it into an embedding space using two alternative strategies: a *CNN layer* or a *flattening layer*. 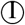 *The CNN layer*. This module processes each frame and projects it into a fixed-dimensional embedding vector of size *out* _*dim*.

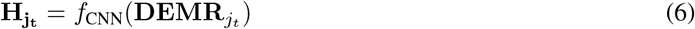

Here, **H** *∈* R^*T ×out*_*dim*^ denotes the output of this module, and *f*_CNN_ represents the CNN layer. 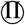 *The flattening layer*. The feature tensor 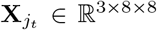 corresponding to a specific frame is directly flattened into a vector of length 192, as follows..

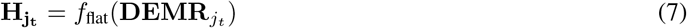

The feature-handling strategies described above were combined with the three frame-sampling strategies (Section 3.2) to generate the overall embedding for subsequent learning. ➀ Time-interval sampling. *T* frames were sampled at different time intervals, and all frames were assigned equal weights. ➁ RMSD-based sampling. *T* frames were sampled based on their RMSD ranges, each corresponding to a single RMSD category (HLC, MLC, or LLC). These frames were weighted according to their RMSD values. ➂ Hybrid-category sampling. *T* frames covering all three RMSD categories (HLC, MLC, and LLC) were sampled, and learnable weights were assigned to them to investigate the respective contributions of different RMSD categories to the prediction tasks. Combining these three scenarios, the weights for the selected *T* frames are listed as below.

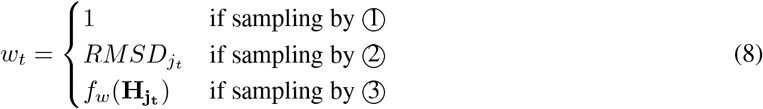

Finally, concatenating (||) the weighted frame-level embeddings yields the overall embedding for further learning.

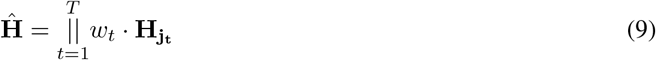

where **Ĥ** denotes the weighted result, which has the same dimensionality as 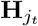.

#### Transformer Encoder

This module was employed to process **Ĥ** and capture complex interactions among different frames, which are essential for learning high-level dynamic representations.

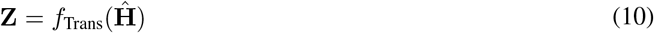

where **Z** represents the output of the Transformer module. *f*_Trans_ denote the functions ofthe Transformer encoder module.

#### Softmax Output

MPP tasks were formulated as a binary classification problem. The final molecular representation **Z** was then passed through a multi-layer perceptron (MLP), and the resulting logits were normalized using the softmax function to produce class probabilities:

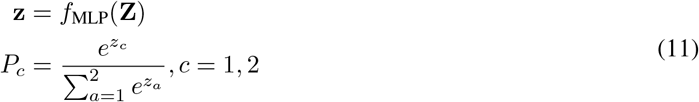

where **z** = [*z*_1_, *z*_2_] *∈* ℝ^1*×*2^ represents the output of the MLP, *z*_1_ and *z*_2_ are the unnormalized logits for the two classes. The function *f*_MLP_ denotes the transformation implemented by the MLP. *P*_*c*_ is the predicted probability of the molecule belonging to class *c* (i.e., positive or negative).

#### Weighted Cross-Entropy Loss

As shown in Table. 1, it is evident that there is a class imbalance between positive and negative samples in MPP tasks. This imbalance can lead to biased predictions and should be carefully addressed. In this study, we incorporated a weighted cross-entropy loss (WCEL) function into the training process. The weights were calculated based on the number of samples in each class, with higher weights assigned to the positive class. This enables the model to focus on and recognize the positive class, thereby enhancing overall classification performance.

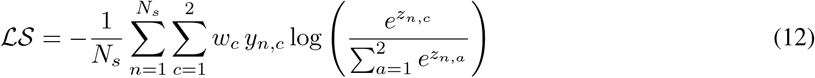

where *ℒS* denotes the loss function, *N*_*s*_ is the number of samples, and *y* represents the ground truth after one-hot encoding; *y*_*n*,*c*_ is the label corresponding to the *c*-th class of the *n*-th sample, and *z*_*n*,*c*_ is the model output for the *c*-th class of the *n*-th sample; *w*_*c*_ is the weight of *c*-th class, defined as 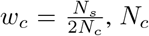, *N*_*c*_ denotes the number of samples belonging to the *c*-th class.

## 4 Experiments and Results

### 4.1 Generated MD datasets

According to the workflow in Fig. 3, all-atom MD simulations were conducted for 8,145 molecules for MPP tasks. These MD simulations were performed on high-performance computing servers equipped with NVIDIA V100 32GB GPUs. All-atom simulations are computationally intensive; as an example, a 10-nanosecond simulation of molecule *TOX22632*, which contains 50 heavy atoms, cost approximately 23 minutes (1,390 seconds) in our experiments. The molecules in our dataset contain between 4 and 197 atoms, which leads to varying computational costs for simulations.

To ensure reliable MD simulations, we monitored the trends of several metrics (temperature, density, total energy and RMSD) during the MD simulations. The evaluation results for *TOX22632* are displayed in Fig. 5 as an example, where the trends are reasonable, indicating that the system is sufficiently stable. Similar evaluation results for other molecules with different sizes can be found in the Appendix B. Finally, to facilitate public access to our MD data, all datasets have been deposited in *Zenodo* repository at https://doi.org/10.5281/zenodo.15788151.

**Fig. 5.**
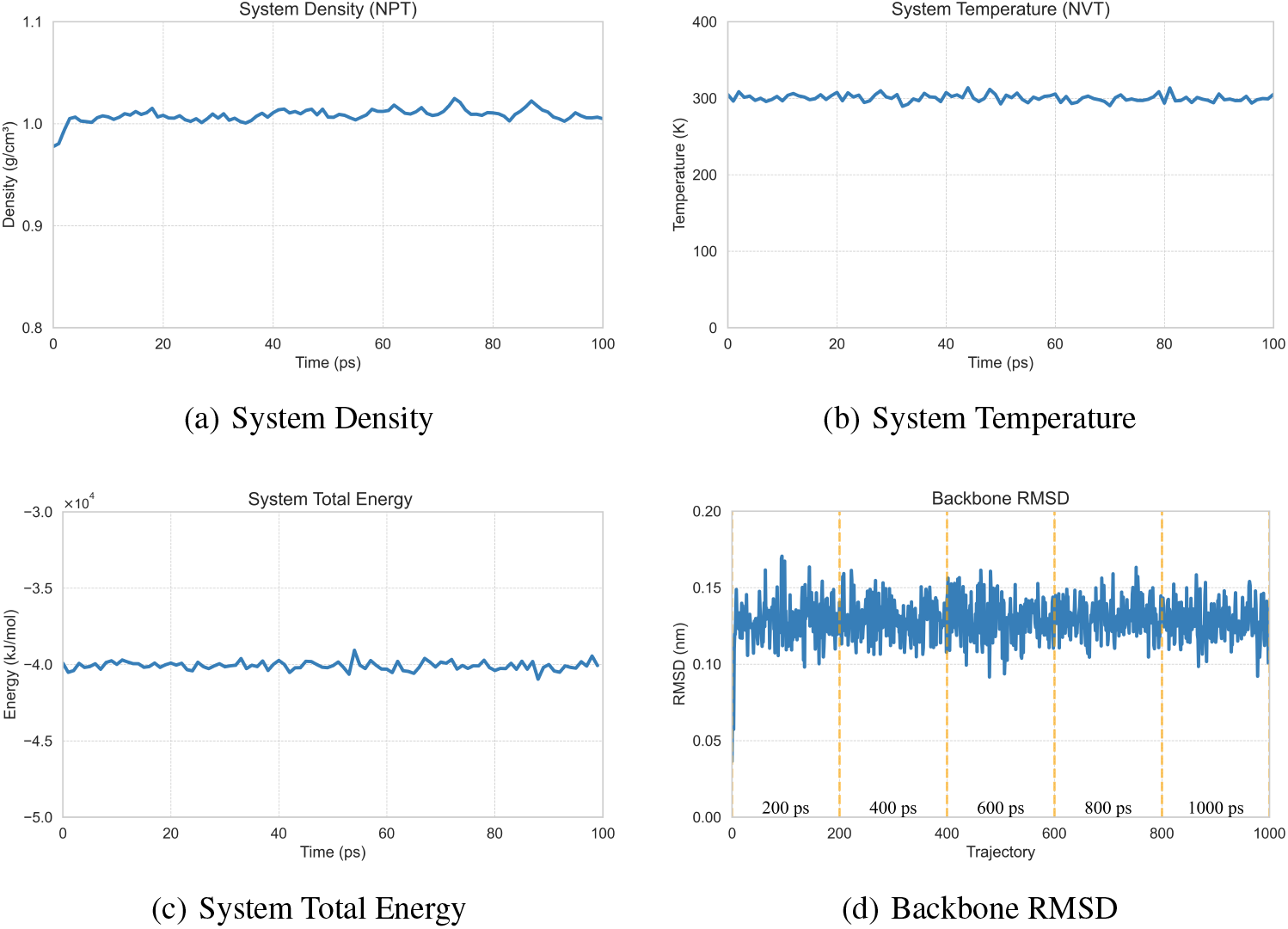
Quality evaluation for the MD simulations on the ‘TOX22632’ system.

### 4.2 Experimental Setting

All the models were trained on NVIDIA RTX 2080Ti GPUs. For each task, the dataset was split into training, validation, and test sets with a ratio of 7 : 1 : 2, using a batch size of 32 and 200 training epochs. To prevent overfitting and enhance generalization, an early stopping strategy with a patience of 10 epochs was employed. Due to the class imbalance, model performance was evaluated using ROC-AUC, Balanced Accuracy (Balanced Acc), and Matthews Correlation Coefficient (MCC).

#### Dynamics-involved learning

The *CNN module* comprises two 2D convolutional layers with 3 *×* 3 kernels. The ReLU activation function is applied after each layer, and dropout is used to prevent overfitting. The CNN output is projected to a 128-dimensional vector (*out* _*dim* = 128) via a fully connected layer. The *Transformer module* consists of 6 encoder layers, and each encoder layer contains 4 self-attention heads. Learnable positional encodings are introduced to capture temporal dependencies more effectively. The classifier has a 2-layer MLP with ReLU activation and Dropout. Final binary classification probabilities are obtained via a *Softmax layer*,facilitating effective discrimination of molecular properties.

#### Baselines

We compared our models with a variety commonly used methods in MPP tasks. ➀ Some of them employ molecular fingerprints (e.g., ECFP and MACCS) as feature representations, and adopt traditional machine learning models for MPP. Four machine learning models were considered, including Decision Tree (DT), KNearest Neighbor (KNN), Logistic Regression (LR), and Naive Bayes (NB). ➁ The *DeepTox* model leverages ECFPs and Deep Neural Networks (DNNs) is often regarded as a modern benchmark for toxicity prediction. Here we also included it in the baseline list. ➂ Graph-based models serve as strong competitors in MPP tasks, and thus we added them as another set of baselines. The molecules were represented as graphs, with the atoms as nodes and the bonds as edges. Basic atomic properties (e.g., atom types) were used to characterize the nodes. Five graph neural network-based models (APPNP, GAT, GCN, GIN, and GraphSAGE) were adopted to learn these molecular graphs and make predictions. ➃ Treating the molecules as SMILES sequences, a pre-trained large language model (ChemBERTa) was employed to extract feature embeddings, which were then fed into a DNN for MPP.

### 4.3 Results and Analysis

#### Overall performance

Table. 2 summarizes the results of both the baselines and our models (Fig. 4). In Table. 2, DEMR ➀, ➁, and ➂ denote the molecular feature representations obtained using sampling strategies ➀, ➁,and ➃, respectively. For clarity, we report the average results for the *toxicity prediction* and *protein-binder identification* tasks. The results demonstrate that our approach achieves superior prediction performance in *toxicity prediction* tasks; however, for *protein-binder identification* tasks, our proposed models (*CNNTransformer* and *Transformer*) for DEMR did not outperform baseline models that rely on traditional static molecular features such as ECFP and MACCS. Possible reasons for the relatively poor performance in *protein-binder identification* tasks are as follows: the three protein-binder identification datasets exhibit extremely low category imbalance ratios; the number of samples in these datasets is relatively less, conditions under which traditional methods often achieve higher accuracy. In contrast, *toxicity prediction* tasks suffer from more severe class imbalance characterized by a low proportion of positive samples and a large overall sample size making them particularly challenging. The proposed DEMR-based model demonstrated superior performance in more severe class imbalance tasks. Notably, using only flattened features (*Transformer* model), rather than CNN-processed features (*CNNTransformer* model), leads to even more promising results.

**Table 2.**
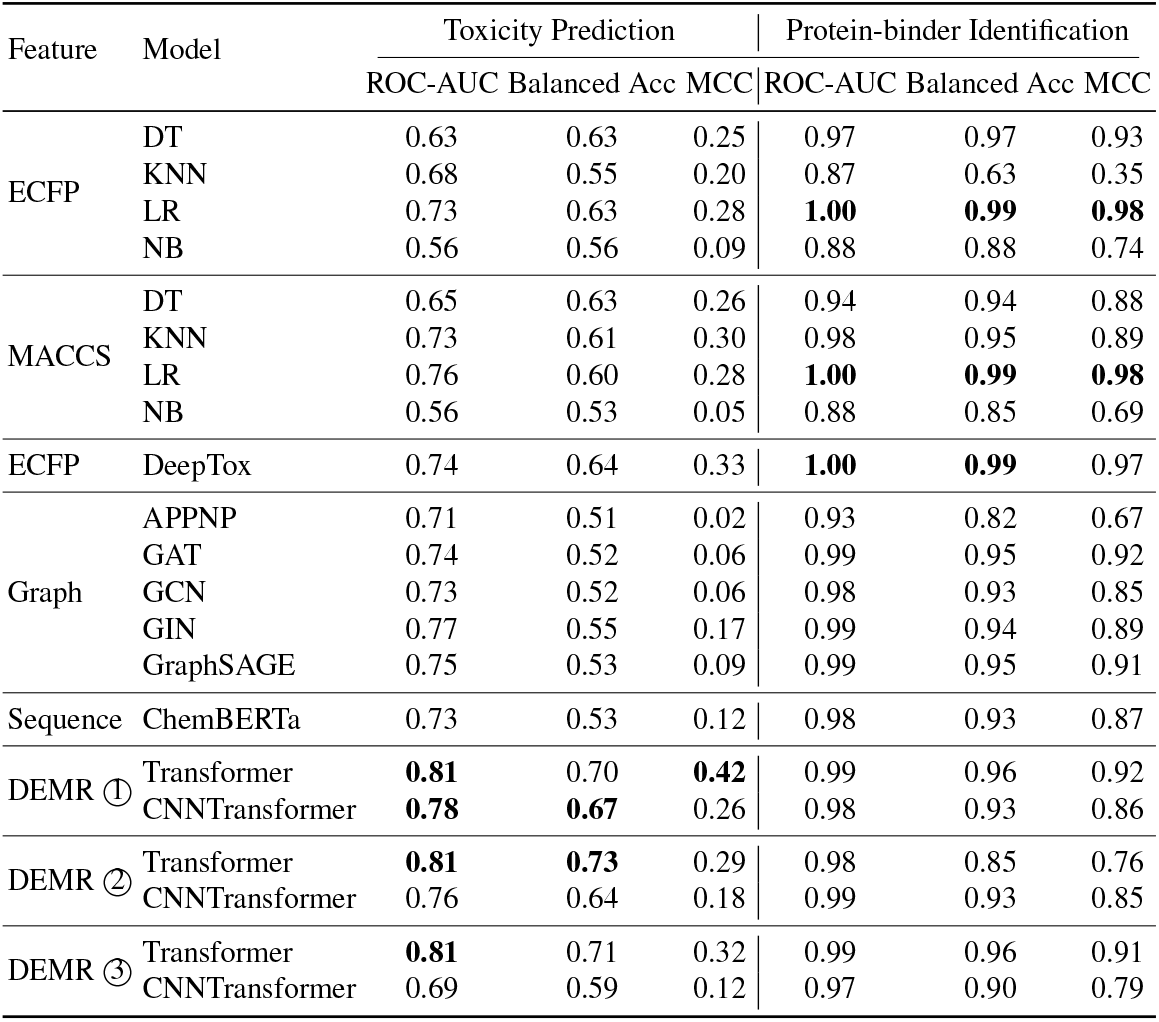
Model performance comparison.

To further demonstrate the effectiveness of our approach, we analyzed the ROC-AUC metrics for all *toxicity prediction* tasks, as shown in Fig. 6. For clearer comparison, several representative baseline models, including LR trained with MACCS and ECFP, and GIN, were evaluated alongside our proposed methods. On average, the *Transformer* model with sampling strategy ➀ or ➂ for DEMR performs the best across most tasks. Moreover, NR-AR-LBD is a well-known challenging task in the field of toxicity prediction [44], and our models based on DEMR achieve competitive results.

**Fig. 6.**
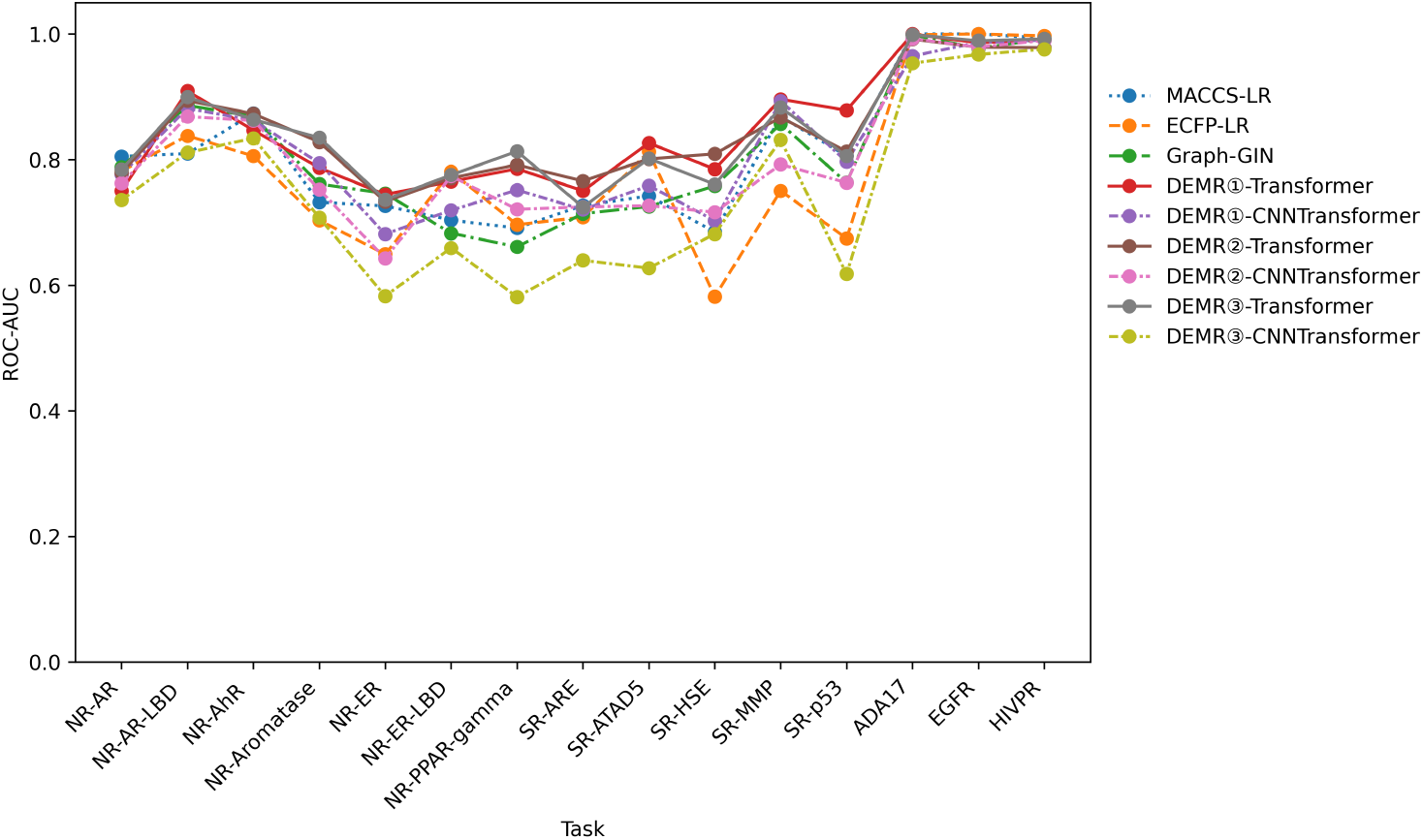
Performance evaluation for specific tasks.

### 4.4 Sampling Strategies Analysis

Here, we analyze the performance of DEMR constructed using three different sampling strategies. More details can be found in Appendix C.

#### Results and analysis for sampling strategies ➀

We sampled the frames at different time intervals *int* for the MPP tasks. For each task, the performance of our models corresponding to different intervals was recorded. The experimental results show that selecting an appropriate sampling frequency for each task could significantly improve the model performance. In contrast, using suboptimal sampling frequencies might significantly impair the performance of the models.

#### Results and analysis for sampling strategies ➁

The sampling strategy ➁ was designed to investigate the effect of molecular conformations on model performance. Models were trained on DEMR of sampled from the HLC, MLC and LLC categories, respectively. The experimental results demonstrate that the model performs relatively better on the DEMR sampled from the HLC and MLC categories. Compared to the LLC category,the HLC and MLC categories exhibit higher RMSDs from the original conformation, implying larger structural variations of molecules. It suggests that introducing extra information on conformational variations into the modeling process will enhance the learning performance of the models.

#### Results and analysis for sampling strategies ➂

Sampling strategy ➂ aims to examine how the model handles different types of molecular conformations (i.e., HLC, MLC, and LLC categories). By monitoring the training process, we conducted statistical and analytical evaluations for the learned weights to assess whether the model is more sensitive to specific RMSD ranges (HLC, MLC, and LLC). The average weights of these three RMSD ranges are shown in Table. 3. Again, the results verify that the model tends to assign higher weights to DEMR sampled in the HLC and MLC categories. This indicates that the molecular conformations with larger structural variations from the original conformation in the MD simulation process carry more information. In contrast, LLC conformations received negative weights, showing that these conformations contribute less to the MPP tasks.

**Table 3.**
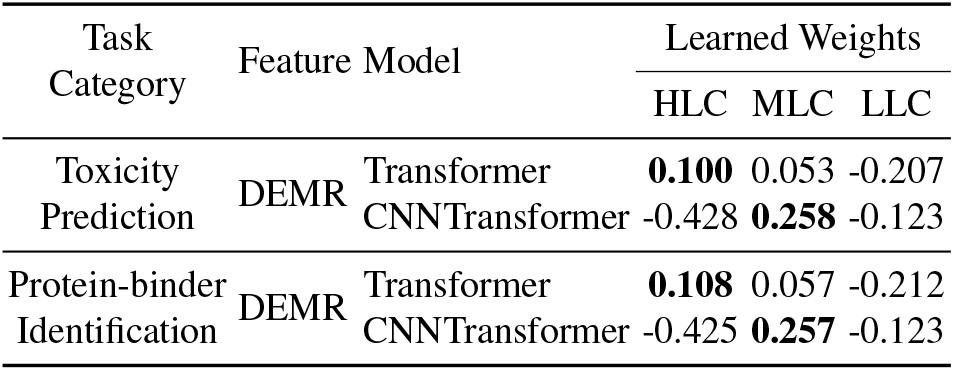
The results of sampling strategies ➂.

### 4.5 Time Cost Analysis

Based on the experimental results of the above three strategies, we found that our models achieved high predictive accuracy using only a few frames (e.g., 10 frames) in nearly half of the tasks. For example, sampled frames resulted in very good predictive performance in the NR-AhR, NR-Aromatase, NR-PPAR-gamma, SR-ATAD5, SR-MMP, ADA17, and EGFR tasks. This finding suggests that the DEMR representation is capable of retaining essential temporal-spatial patterns even with sparsely sampled frames, thereby achieving a favorable trade-off between computational efficiency and predictive performance. The total training time for such a configuration was approximately 2, 000*s* on an NVIDIA RTX 2080Ti GPU. Given the rapid advancement of GPU architectures and processor technologies, such computational costs are considered acceptable for handling large-scale molecular datasets in MPP studies. Moreover, using more advanced GPUs in the future will reduce such costs further and support dynamics-involved studies in MPP tasks even more.

## 5 Conclusion

In this paper, we conducted comprehensive MD simulations on several representative MPP datasets and established a series of large-scale molecular dynamics datasets specifically designed for MPP tasks. These curated datasets occupy approximately 200 GB of storage and are publicly available, providing a valuable benchmark resource for future dynamics-driven molecular modeling research. This contribution not only enriches the available data foundation for MPP studies but also opens up new possibilities for exploring temporal and conformational information in molecular representations. Based on such data, we proposed the molecular representation, DEMR, and two deep learning methods for dynamics-based MPP tasks. Extensive experiments and analyses demonstrated that our approach achieved better performance for MPP tasks with severely imbalanced classes. Moreover, the experimental results showed that a properly configured sampling frequency can effectively enhance the representation ability of DEMR and greatly improve the model performance in MPP tasks. On the other hand, DEMR sampled in the HLC categories are often more advantageous in guiding an accurate MPP model.

## Acknowledgments

This work is supported by Hong Kong Metropolitan University (Project PFDS/2024/01), Hong Kong Research Grants Council (Project UGC/FDS16/E16/23), and the National Natural Science Foundation of China (Projects 62376161 and U24A20322).

## Appendix A CHARMM General Force Field

The CHARMM (Chemistry at Harvard Macromolecular Mechanics) General force field is a widely used classical force field developed originally at Harvard University. It was designed to simulate the structural and dynamic behaviors of biomolecules including proteins, nucleic acids, lipids, and small molecules. The total energy function *E*_total_ included bonded terms and non-bonded terms, and it could be expressed as:

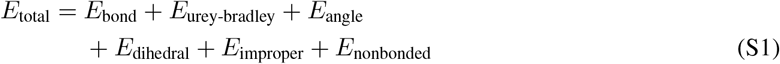

where *E*_bond_, *E*_urey-bradley_, *E*_angle_, *E*_dihedral_, *E*_improper_ and *E*_nonbonded_ represent the energy of bond stretching, Urey-Bradley, angle bending, dihedral torsion, improper torsion, and nonbonded(van der Waals and Electrostatics), respectively. In this study, we used *CGenFF* to generate the topologies of molecules through its online interface (https://cgenff.com/).

The CHARMM force field defines an extensive set of atom types to represent atomic behavior in diverse chemical environments. These atom types are characterized by elemental identity (e.g., *C, H, O, N, S*,*P*, etc), hybridization state (e.g., sp^3^, sp^2^, aromatic), and bonding context. Each atom type is associated with specific parameters, such as partial charges, van der Waals radii, bond lengths, bond angles, and dihedral terms. A complete description could be found in the *CGenFF* documentation.

In our experiments, *CGenFF* was unable to process some non-standard molecular structures, resulting in parameterization failures for some molecules. Common reasons for these failures include unsupported metal complexes (e.g., *Zn, Fe*); the presence of radicals or NO-containing groups that complicate electron counting; undefined atom types for elements such as *B, Si*, or phosphoryl groups (*P=O*); and structural issues such as incomplete molecules (e.g., unclosed rings or dangling atoms) or incorrect atomic charges. These issues prevented *CGenFF* from correctly recognizing the chemical environment of molecules, and some molecules could not be successfully processed. Consequently, the number of molecules in our constructed MD dataset differed from the original dataset. Nonetheless, this did not diminish the significance of our efforts in constructing a high-quality MD dataset.

## B Quality evaluation for Our MD Simulations

We evaluated the quality of MD simulations for some representative molecules, as shown in Fig.S1. For each dataset, two representative molecules were selected and presented as examples. The results confirmed that all simulated systems satisfied the expected physical conditions:

1. System density remained close to 1*g/cm*^3^;
2. System temperature was maintained at 300*K*;
3. The total energy of the system remained stable throughout the simulation;
4. The backbone RMSD values of the MD trajectories were relatively low.

**Fig. S1.**
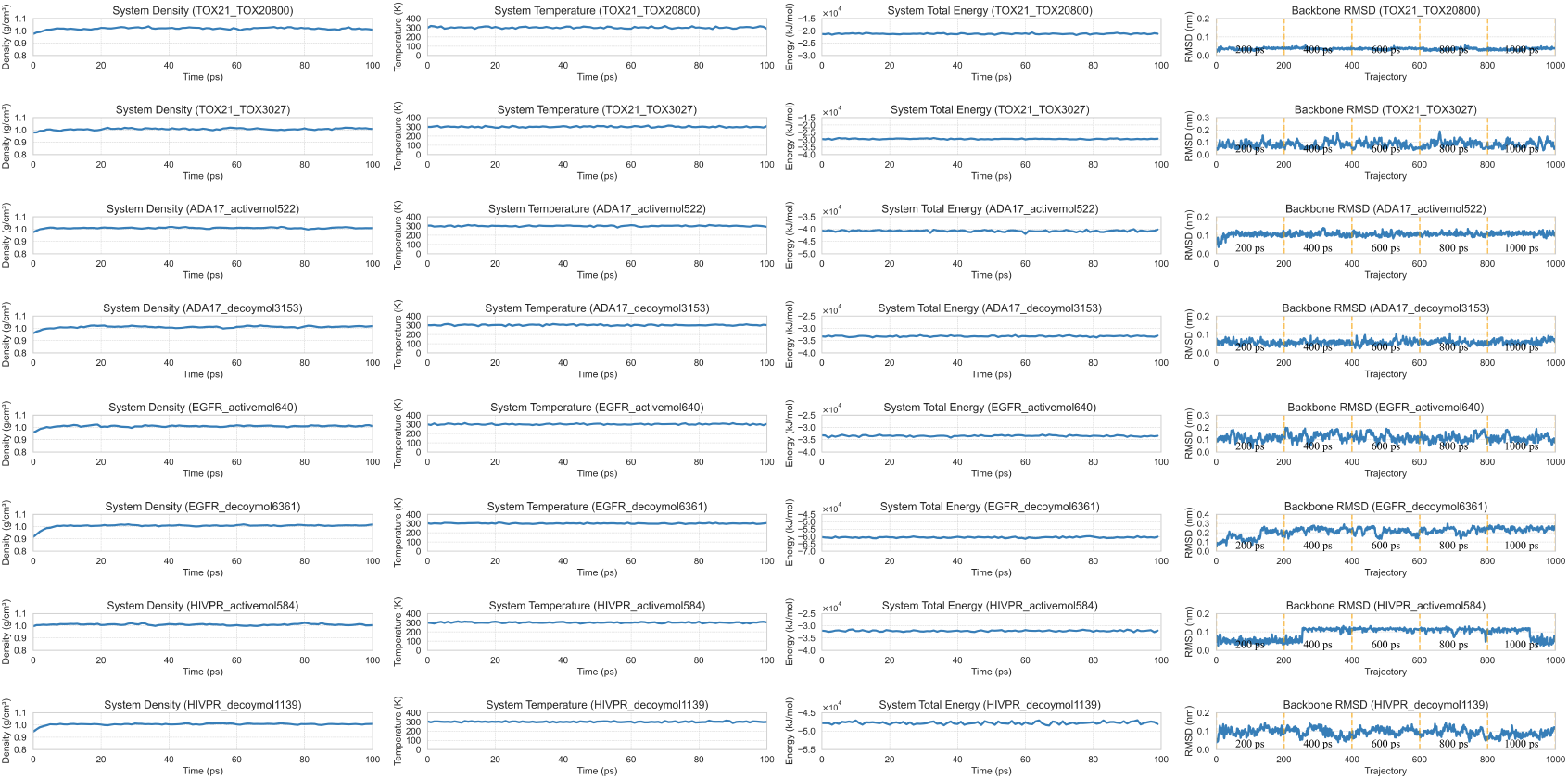
Quality evaluation for the MD simulations on different datasets.

The quality evaluation results indicated that the molecular structures remained structurally stable and well equilibrated throughout the MD simulations.

Table. S1 summarizes the molecular system sizes information and the MD simulation cost time. With increasing molecular size, MD simulations demanded significantly more time and computational resources, primarily due to the larger number of atoms and interactions involved.

**Table S1.**
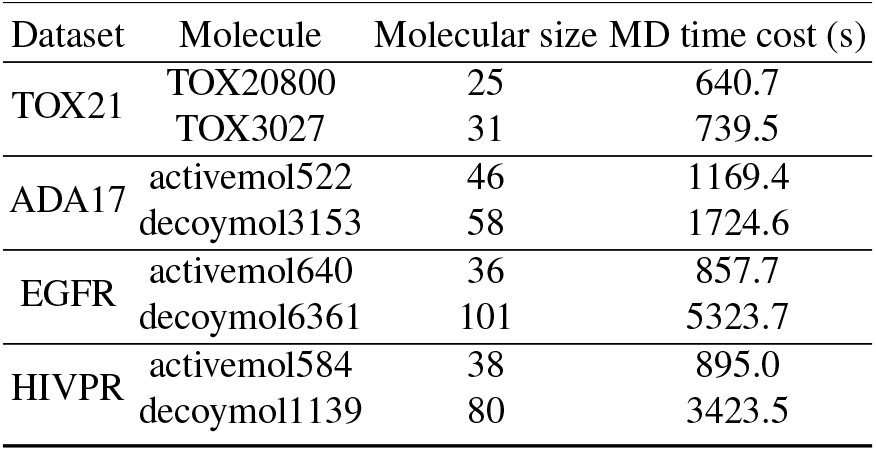
Molecular information in MD simulations.

| Dataset | Molecule | Molecular size | MD time cost (s) |
| --- | --- | --- | --- |
| TOX21 | TOX20800 | 25 | 640.7 |
|  | TOX3027 | 31 | 739.5 |
| ADA17 | activemol522 | 46 | 1169.4 |
|  | decoymol3153 | 58 | 1724.6 |
| EGFR | activemol640 | 36 | 857.7 |
|  | decoymol6361 | 101 | 5323.7 |
| HIVPR | activemol584 | 38 | 895.0 |
|  | decoymol1139 | 80 | 3423.5 |

## C Specific analysis for sampling strategies

### Analysis for sampling strategies ➀

We analyzed the impact of model performance in sampling strategy 1. Fig. S2 presented the average ROC-AUC results of *Transformer* and *CNNTransformer* of sampling strategy 1, respectively. Specifically, our results indicated that selecting an appropriate sampling frequency for each task could significantly improve the performance of both models. In contrast, using suboptimal sampling frequencies might significantly impair the performance of the models. These findings highlight the importance of task-specific sampling frequency selection for the optimal exploitation of DEMR features.

**Fig. S2.**
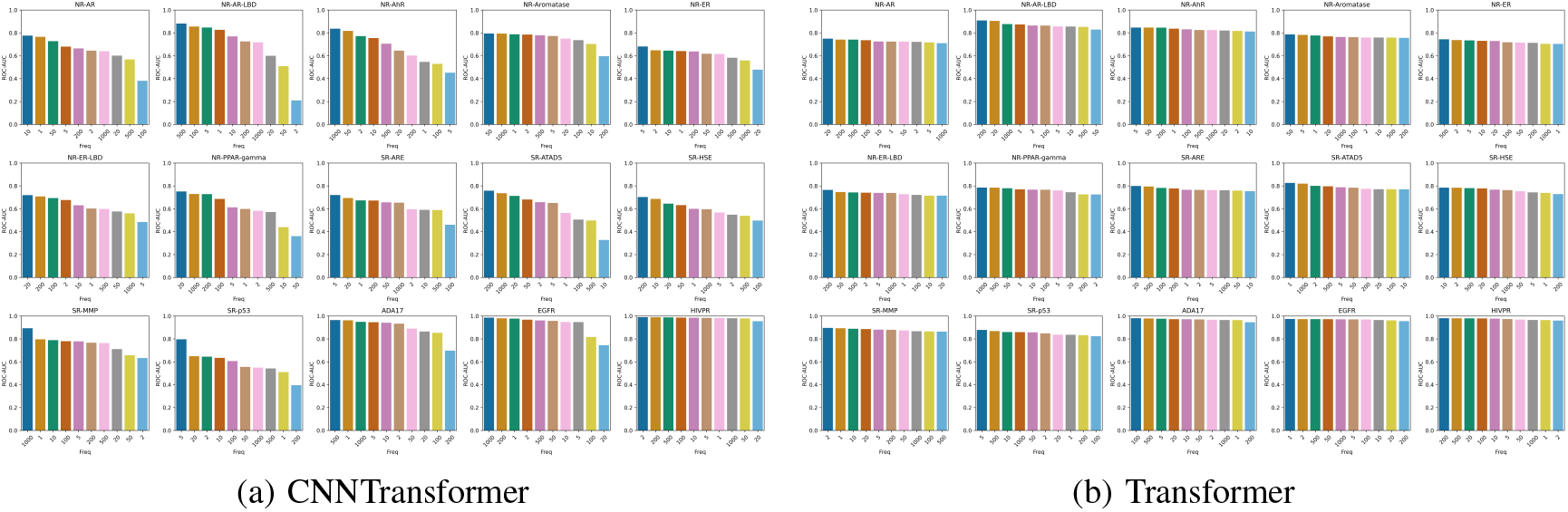
ROC-AUC results in sampling strategy ➀ for different models.

### Analysis for sampling strategies ➁

The sampling strategy ➁ was designed to investigate the effect of molecular conformations on model performance. Models were trained on DEMR of HLC, MLC and LLC category, respectively. Fig. S3 shows the ROC-AUC results of the different RMSD ranges (HLC, MLC and LLC) DEMR under the *Transformer* and *CNNTransformer*, respectively. Overall, under a fixed sampling frequency, the *Transformer* showed consistent performance across most tasks, demonstrating limited sensitivity to HLC, MLC, and LLC DEMR. In contrast, the *CNNTransformer* achieved significantly better performance on the HLC and MLC DEMR compared with the LLC DEMR. This suggests that the HLC and MLC DEMR are more beneficial for the *CNNTransformer*, potentially due to its stronger reliance on spatial and temporal dynamics. Especially in the SR-MPP and HIVPR datasets, models trained only based on the LLC DEMR show significant performance degradation. We attributed this to the insufficient structural diversity of LLC DEMR, which might fail to capture potentially bioactive or functionally relevant molecular conformations. This limitation in LLC DEMR hindered the ability of the model to learn essential patterns. However, while HLC DEMR provided richer structural diversity, they might include extreme molecular conformational.

**Fig. S3.**
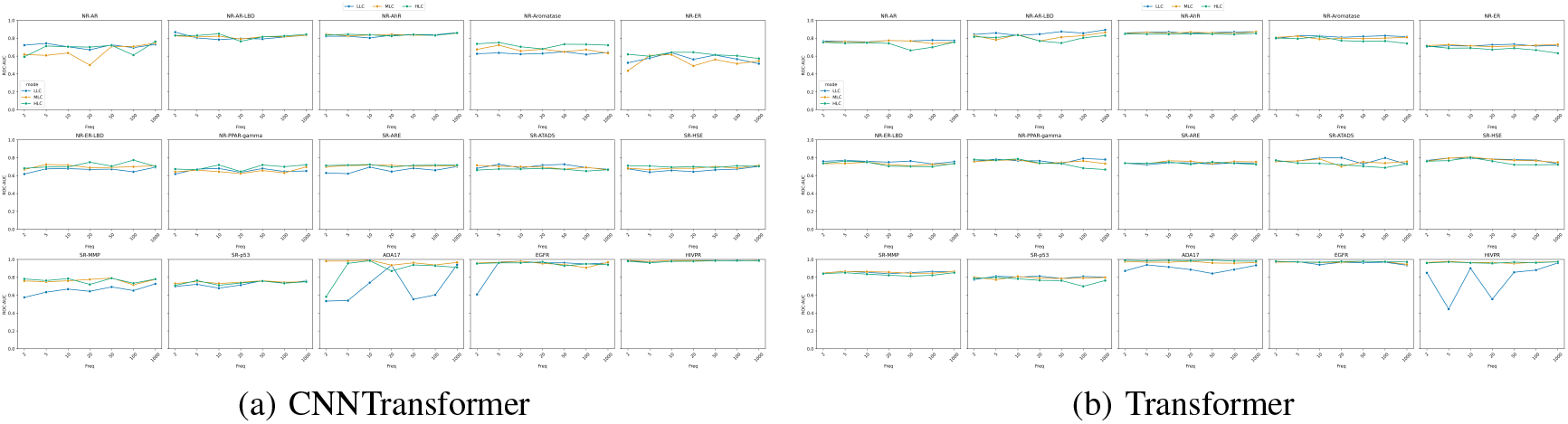
ROC-AUC results in sampling strategy ➁ for different models.

